# Climatic warming strengthens a positive feedback between alpine shrubs and fire

**DOI:** 10.1101/043919

**Authors:** James S Camac, Richard J Williams, Carl-Henrik Wahren, Ary A Hoffmann, Peter A Vesk

**Author notes:** J.C. conceived, designed and performed the experiments, field surveys and analysis; C-H.W, R.W, A.H and P.V. supervised the development of this work, aided in data collection and provided statistical advice. All authors contributed to the writing of this manuscript. The authors declare that they have no competing financial interests.

## Abstract

Climate change is expected to increase fire activity and woody plant encroachment in arctic and alpine landscapes. However, the extent to which these increases interact to affect the structure, function and composition of alpine ecosystems is largely unknown. Here we use field surveys and experimental manipulations to examine how warming and fire affect recruitment, seedling growth and seedling survival in four dominant Australian alpine shrubs. We found that fire increased establishment of shrub seedlings by as much as 33–fold. Experimental warming also doubled growth rates of tall shrub seedlings and could potentially increase their survival. By contrast, warming had no effect on shrub recruitment, post-fire tussock regeneration, or how tussock grass affected shrub seedling growth and survival. These findings indicate that warming, coupled with more frequent or severe fires, will likely result in an increase in the cover and abundance of evergreen shrubs. Given shrubs are one of the most flammable components in alpine and tundra environments, warming is likely to strengthen an existing feedback between woody species abundance and fire in these ecosystems.

## Introduction

Accurately forecasting the effects of climatic warming on vegetation dynamics requires an understanding of the mechanisms by which climate and vegetation interact. Most forecasting models include the direct effects of climatic conditions on species distributions, but largely ignore disturbances, particularly their type, frequency and severity (Guisan & Thuiller, 2005; Tucker *et al*., 2012). Recurrent disturbance is integral to all ecosystems, strongly influencing fundamental demographic processes such as recruitment and mortality, and thus the composition and structure of plant communities and biomes worldwide (Bond *et al*., 2005; Turner, 2010). Disturbance regimes are already changing as a consequence of climate change (Turner, 2010; Westerling *et al*., 2011; Bradstock *et al*., 2014). It is therefore imperative we understand how the effects of disturbance will interact with climate change, and whether such effects amplify or diminish how vegetation responds to changing temperature and moisture (Post & Pedersen, 2008; Camac *et al*., 2015; Enright *et al*., 2015)

Understanding the feedback between climate, vegetation and fire will be particularly important in order to accurately predict the trajectory of ecosystem change in coming decades (Bowman *et al*., 2009). In addition to the direct effects of warming on fire weather, changing climate may also indirectly alter fire regimes by altering vegetation productivity, structure and composition (Keeley *et al*., 2012; Matthews *et al*., 2012; Bowman *et al*., 2014). Such feedbacks may be positive or negative, and have been documented in tropical savanna (Hoffmann, 2003), boreal forests (Krawchuk & Cumming, 2011), grasslands (Flannigan *et al*., 2009) and alpine and arctic environments (Goetz *et al*., 2007; Wookey *et al*., 2009).

Alpine and arctic vegetation are considered to be particularly vulnerable to the effects of changing climate (Engler *et al*., 2011; Dullinger *et al*., 2012; Elmendorf *et al*., 2012a). In these ecosystems, field manipulative experiments have shown that climate directly influences plant phenology (Hoffmann *et al*., 2010; Dorji *et al*., 2013), reproduction (Klady *et al*., 2011), morphology (Hudson & Henry, 2009), growth (Hollister *et al*., 2005), floristic composition (Elmendorf *et al*., 2012a) and biotic interactions (Klanderud, 2005). However, most studies in these ecosystems have focused on mature plant responses in undisturbed vegetation (Briceño *et al*., 2015). Few have included disturbance as a factor (but see: Munier *et al*., 2010; Graae *et al*., 2010; Camac *et al*., 2015), or examined the influence of climate change on seedlings in post-disturbance environments. As a consequence, little is known about how climate affect seedling demographic rates (Briceño *et al*., 2015), particularly in alpine and tundra post-disturbance conditions. This is despite mounting evidence that seedling regeneration is important in alpine and arctic ecosystems (Venn & Morgan, 2009; Briceño *et al*., 2015), particularly for woody species (Camac *et al*., 2013; Williams *et al*., 2014). Seedlings are the life stage that determines the long-term persistence of a species as well as its capacity to establish in new areas (Walck *et al*., 2011). As such, in order to accurately predict future trajectories of vegetation change in alpine and arctic ecosystems, it is impertative we understand how seedlings respond to both changing climate and disturbance regimes (Walck *et al*., 2011; Briceño *et al*., 2015).

## Feedback between climatic warming, fire and shrubs

In alpine and arctic ecosystems, warming experiments and long-term monitoring have documented significant increases in the growth and cover of woody species (Sturm *et al*., 2001; Walker *et al*., 2006; Myers-Smith *et al*., 2011). The frequency and extent of wildfires in these environments have also increased over recent decades, a trend expected to continue (Westerling *et al*., 2006; Flannigan *et al*., 2009; Qiu, 2009; Bradstock *et al*., 2014). Current evidence from these ecosystems indicate that shrub recruitment and encroachment is highest in disturbed areas and lowest in areas with minimal bare ground cover (Williams & Ashton, 1987; Batllori *et al*., 2009; Frost *et al*., 2013). Evidence also suggests that climatic warming is likely to increase growth rates of woody species (Arft *et al*., 1999; Elmendorf *et al*., 2012b; Myers-Smith *et al*., 2011) and that shrubs are potentially the most flammable vegetation component in these ecosystems (Williams *et al*., 2006; Higuera *et al*., 2009; Fraser *et al*., 2016). Thus, increases in the cover of woody species in alpine and arctic environments may increase the flammability of these ecosystems.

The results of these studies suggest a positive feedback could exist between warmer temperatures, woody species and fire in alpine and tundra environments (Fig. 1a). Specifically, warmer temperatures may lead to more frequent and severe fire, which in turn, may increase recruitment opportunities (i.e. more bare ground) for woody species and increase shrub thickening both within and beyond shrub boundaries (Racine *et al*., 2004; Lantz *et al*., 2013). If this effect is coupled with an increase in the growth and survival of shrub seedlings, highly flammable fuels will accumulate at a faster rate, have higher landscape connectivity, and ultimately lead to increases in the likelihood of fire. The consequence of which will further increase shrub recruitment opportunities. Thus, warming could strengthen an existing climate-disturbance feedback, that not only has the potential to cause rapid changes in the composition and structure of alpine and arctic vegetation, but also has serious social, biodiversity and carbon sequestration consequences (Mack *et al*., 2011).

**Fig. 1.**
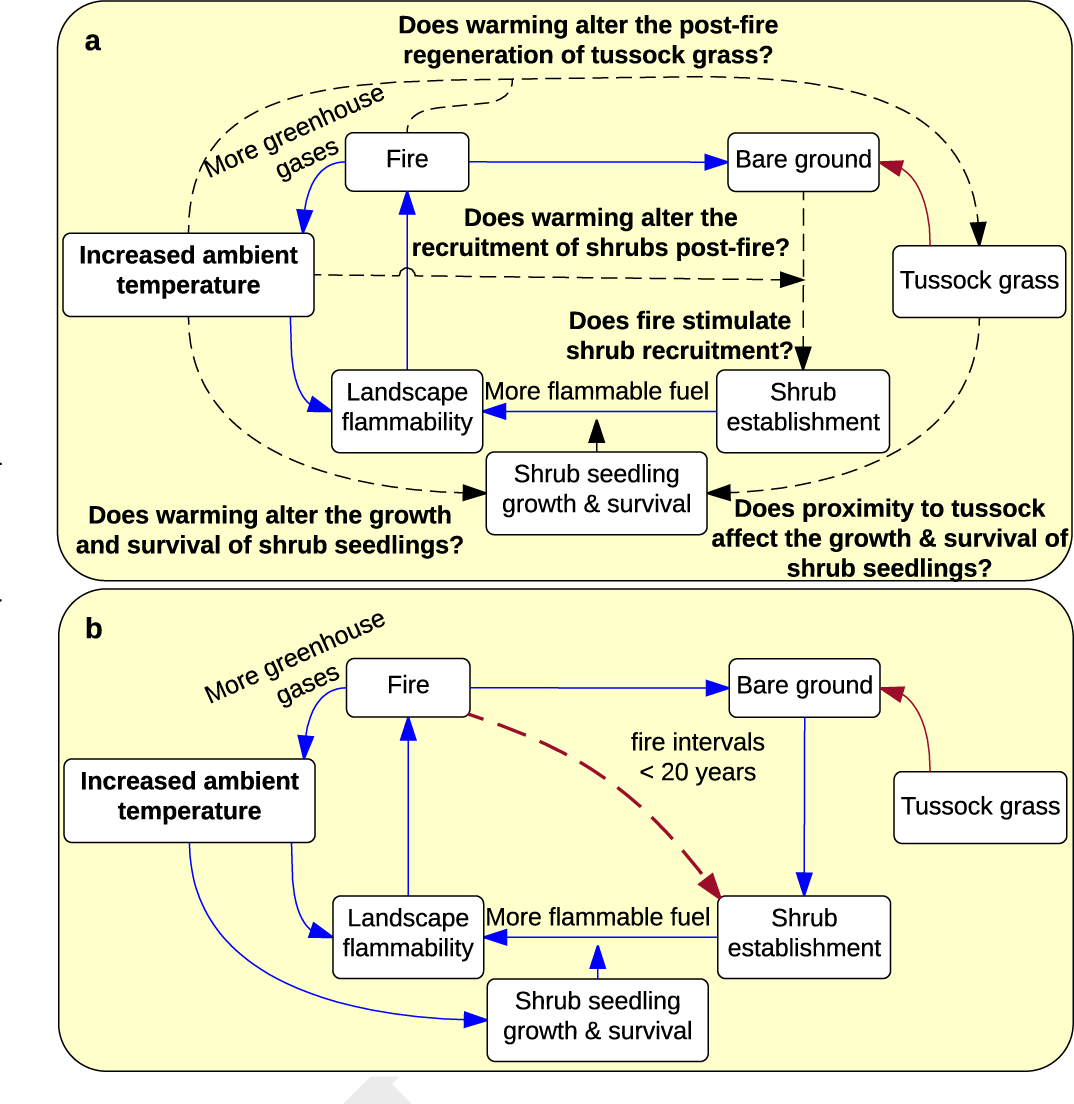
Conceptual diagram illustrating positive warming-shrub-fire feedback. (a) Hypothesised positive feedback loop between fire, climatic warming and landscape flammability. Solid lines = known mechansims; Dashed lines = mechansims that may exacerbate or diminish this feedback but which we have a paucity of information on. (b) Feedback updated based on new evidence obtained from experimental and field surveys conducted in this study. Also includes an example of a potential, but unlikely, effect (red dashed line) that could break this feedback. Blue = positive relationships, Red = negative relationships.

While paleoecological studies have indicated that such a feedback may exist in the arctic (Higuera *et al*., 2008, 2009), there is a paucity of information on what may strengthen, mitigate or break this feedback. For example, we have little information on whether fire will stimulate shrub seedling recruitment in alpine or tundra ecosystems, nor do we know how rates of seedling growth and survival will be affected under warmer, more exposed, post-fire conditions. Furthermore, we do not know how tussock grasslands will respond to warmer post-fire conditions, how grasses affect vital rates of shrub seedlings, or whether such effects are altered by warmer post-fire conditions.

Here, we examine these unknowns and their impacts on this proposed feedback between climatic warming, shrubs and fire in Australian alpine vegetation. As with other tundra ecosystems, Australian alpine landscapes consist of a mosaic of vegetation types dominated by either herbaceous or woody species (Williams *et al*., 2014). They have also experienced significant changes in climate. Since 1979, mean growing season temperatures have risen by approximately 0.4°C and annual precipitation has fallen by 6% (Wahren *et al*., 2013), with a consequent decline in snow pack depth (Sánchez-Bayo & Green, 2013). These climatic changes have been correlated with a 10 to 20% increase in shrub cover and a 25% decline in graminoids cover (Wahren *et al*., 2013). Much of the Australian Alps has also been burnt by recent (2003 and 2006) wildfires, the frequency and severity of which are expected to increase as a consequence of further climate change (Bradstock *et al*., 2014; Williams *et al*., 2014).

In this study we took advantage of recent fires in alpine open heathland, a plant community that occupies ca. 25% of the Australian alpine landscape (Williams *et al*., 2014). Under global warming, open heathland is likely to encroach upon grasslands, wetlands and herbfields (including the nationally endangered snow patch herbfields; Williams *et al*., 2015), and is itself susceptible to shrub thickening McDougall, 2003). To identify some of the biotic and abiotic factors that affect shrub establishment and how warmer post-fire conditions affect shrub seedling vital rates, we combined field observations of post-fire seedling densities with a warming experiment that used seedlings of four dominant Australian evergreen fire-killed shrubs: *Grevillea australis* (Proteaceae; a tall shrub), *Asterolasia trymalioides* (Rutaceae; a prostrate shrub), *Phebalium squamulosum* (Rutaceae; a tall shrub) and *Prostanthera cuneata* (Lamiaceae; a tall shrub).

This combination of data allowed us to quantify the following unknowns in the hypothesised climate-fire-shrub feedback loop (Fig. 1a):

1. Does fire stimulate shrub recruitment?
2. Does warming alter the recruitment of shrubs post-fire?
3. Does warming alter the growth and survival of shrub seedlings?
4. Does proximity to tussock grass affect the growth and survival of shrub seedlings?
5. Does warming alter the post-fire regeneration of tussock grass?

## Methods

Below, we provide a summary of the experimental design and analysis. Full details are presented in the Supporting Information.

### Study Sites

The Australian Alps are an ideal place to examine the proposed fire-climate-vegetation feedback because they have all the elements needed to demonstrate such a feedback. The Alps are subject to recurrent landscape fire (approximately every 50 to 100 years), and were burnt extensively in 2003 (Williams *et al*., 2006, 2014). The vegetation consists of a range of life forms (graminoids, forbs and shrubs), of which shrub abundance strongly determines landscape flammability (Williams *et al*., 2006; Fraser *et al*., 2016). There is also evidence that both fire regimes (Bradstock *et al*., 2014) and the relative abundance of life forms in the Australian Alps have changed as a consequence of recent climate change (Wahren *et al*., 2013; Camac *et al*., 2015).

We investigated shrub-grass dynamics in open heathland on the Bogong High Plains, Australia. Open heathland is a common Australian alpine vegetation type, occupying approximately 25% of the treeless landscapes above 1600m in the Australian Alps. It is an ecotone between closed heathland (>70% shrub cover) and tussock grassland (shrub cover < 20%). Open heathland is dominated by the shrub *Grevillea australis* (shrub cover generally 20–60%) with inter-shrub regions occupied by snow grasses (*Poa hiemata* and *Poa costiniana*) and other herbaceous species. *Grevillea australis* is an obligate seeding shrub, and establishment of seedlings is dependent on disturbance creating bare ground in the grass–herb sward (Williams & Ashton, 1987). Relative to tussock grassland, open heathland is a highly flammable plant community (Williams *et al*., 2006; Fraser *et al*., 2016). A consequence of this differential flammability led to the 2003 wildfires on the Bogong High Plains burning approximately 60% of open heathland and only 13% of tussock grasslands. Following these fires, seedling regeneration of *Grevillea australis* and other shrubs was prolific (Williams *et al*., 2014).

### Open Top Chamber Experiment

In March 2010, at 1750 m a.s.l, we burnt 32 randomly selected mature (60 cm tall and 1.5 m2) *Grevillea australis* shrubs in an open heathland site that was not burnt by wildfire in 2003 or 2006. This created patches of bare ground approximately 0.7 m2 surrounded by burnt tussock grass, simulating disturbance of individual shrubs in open heathland burnt by wildfire. We collected seedlings of dominant alpine shrub species from a nearby (<2 km) site of similar altitude burnt by 2006 wildfires. We collected seedlings of two dominant open heathland shrub species *Grevillea australis* (Proteaceae; a tall shrub) and *Asterolasia trymalioides* (Rutaceae; a prostrate shrub), a dominant closed heathland species *Prostanthera cuneata* (Lamiaceae; a tall shrub) that typically grows on warmer aspects and a species common to both open and closed heathland *Phebalium squamulosum* (Rutaceae; a tall shrub). We focused on these four species because they are common in the Australian Alps; are fire-killed and thereby re-establish via seed (the dominant shrub post-fire strategy in the Australian mainland Alps; Walsh & McDougall, 2004); and under climatic warming, have the potential to increase in cover and height within heathlands, and invade non-shrubby plant communities such as alpine grasslands and herbfields.

A total of 640 seedlings, 256 *Grevillea* (half used in *Poa* inter-tussock experiment—see below) and 128 for each of *Asterolasia, Prostanthera* and *Phebalium* were used. Four seedlings per species were randomly selected and transplanted into a 4×4 square grid in the center of each burnt patch, with 14cm between individuals and the edge of the patch, which was dominated by resprouting tussock grass *Poa hiemata*. To examine interactions between tussock grass and shrub seedlings we also randomly transplanted four additional *Grevillea australis* seedlings into various sized bare gaps between burnt *Poa hiemata* tussocks that were immediately surrounding the 4×4 bare ground square grid (Fig. S1). The experimental site was fenced to prevent grazing by deer and horses. We detected no obvious signs of herbivory by invertebrates or hares within our plots.

To simulate near-term warmer conditions indicated by the IPCC (2013), we randomly assigned Open Top Chambers (OTCs) to half (16) the plots, with the remainder treated as unwarmed controls. The chambers were constructed following the International Tundra Experiment (ITEX) protocols (Molau & Mølgaard, 1996). OTCs were placed over plots, ensuring all seedlings (including inter-tussock shrub seedlings) occurred within the 1.1 m^2^ open top to minimise edge effects. OTCs were placed out at the start of the growing season (October) where they remained until first snowfall (early June). This procedure was repeated for six growing seasons from May 2010 to May 2016.

Microclimatic conditions were measured hourly using Onset Micro Stations (Onset Computer Corporation, Bourne, MA, USA) at four control and four OTC plots. Across six growing seasons (1281 growing season days), OTCs simulated warmer conditions at the lower end of IPCC (2013) projections (Fig. S6-S8). OTCs passively increased average ambient and soil temperatures by 0.9°C. Minimum and maximum temperatures were also raised in both ambient air (min: 1.1°C; max: 2°C) and soil (min: 2°C; max: 0.1°C). Chambers only marginally decreased soil moisture by 0.1% and relative humidity by 0.7%.

Seedling survival, maximum height and stem diameter (nearest mm measured with Vernier calipers) were initially recorded in May 2010 and then subsequently re-measured at the end of each growing season (May-June). At the same time, we recorded the distance to the nearest tussock or grass seedling in each of four cardinal directions for *Grevillea* seedlings growing in inter-tussock gaps. Thus, temporal changes in inter-tussock gap size could be due to either vegetative growth of resprouting tussocks, or the establishment of grass seedlings. We did not measure individual characteristics (e.g. height and basal diameter) of surrounding tussock grass because we could not distinguish individuals, and because height varied substantially throughout the season. Numbers of natural *Grevillea australis* recruits establishing within the plots were also recorded and identified for the first two years of the experiment.

### Seedling gradient study

We used 40 open heathland monitoring sites established after the 2003 fires (Williams *et al*., 2006; Camac *et al*., 2013). These sites consisted of 30 burnt sites and 10 sites known to be unburnt for over 70 years. The disparity in sample number between burnt and unburnt treatments was due to very few open heathland sites of sufficient size (0.25 ha) remaining unburnt after both the 2003 and 2006 fires. In the summer of 2011-12, at each site, seedling density/m^2^ was estimated using 40 quadrats, each 1 m^2^, that were evenly distributed in groups of 10 along four 50 m transects, with 10 m between transect lines, subsampling an area of 2000 m^2^. Within plots we recorded the number and maximum height of *Grevillea* and *Asterolasia* seedlings. For unburnt sites we counted the number of mature *Grevillea* plants (>0.5 m^2^) within 5 m of each transect. In burnt sites, this required counting the number of skeletons (there were no living adults at any burnt site) that still persisted post-fire. We were unable to estimate numbers of adult *Asterolasia* because this species does not have a persistent woody skeleton post-fire. Site level data, elevation and Topographic Wetness Index (TWI; a measure of plant available water, Moore *et al*., 1993) were obtained from a 30 m resolution digital elevation model. Lastly, for burnt sites, fire severity was estimated by twig diameters (Whight & Bradstock, 1999), collected immediately after the 2003 fires (Williams *et al*., 2006).

### Data analysis

We built multiple hierarchical models to examine how increased temperature and other factors influenced shrub seedling recruitment, growth and mortality, as well as tussock-grass gap dynamics. For each model we used Bayesian inference and fitted models in `R 3.3.1` (R Core Team, 2016) using package `rstan 2.12.1` (Stan Development Team, 2016). Detailed information about experimental design and analysis is provided in the Supporting Information. Data and source code are available at: https://github.com/jscamac/Alpine_Shrub_Experiment. In order to aid in the reproducibility of this work, our code was written using a remake framework (FitzJohn, 2015), such that others can readily reproduce our entire workflow from data processing, through to producing a pdf of this manuscript by calling `remake() in R`.

## Results

### Drivers of shrub seedling establishment

We first investigated how altitude, Topographic Wetness Index (TWI), adult density, fire and fire severity (as measured by post-fire twig diameters—see Supporting Information) influenced the density of *Grevillea* and *Asterolasia* seedlings (the two dominant shrubs of alpine open heathland). Across the 40 alpine sites surveyed in 2011-12 we found that the abundance of *Grevillea* (Fig. 2) and *Asterolasia* (Fig. S2) seedlings was strongly influenced by the occurrence of fire. Sites burnt in 2003 had seedling densities between 15 and 33 times higher than unburnt sites. The mean seedling density of *Grevillea*, was 1.31/m^2^ at burnt sites and 0.04/m^2^ at unburnt sites. *Asterolasia* had similar mean densities: 1.65 and 0.11 seedlings/m^2^ at burnt and unburnt sites, respectively. For both species, seedling density increased with increasing fire severity (i.e. sites with larger post-fire twig diameters). As hypothesised, pre-fire adult density also positively influenced *Grevillea* seedling density. For both shrub species, we detected no change in seedling density along a 190m elevational gradient (equivalent to a 1.5°C change in mean temperature; Brown & Millner, 1989). This is consistent with the field warming experiment (see below) which indicated rates of recruitment (Fig. S3) and mortality in *Grevillea* is largely insensitive to a 0.9°C change in temperature. We detected no strong effect of Topographic Wetness Index for either species.

**Fig. 2.**
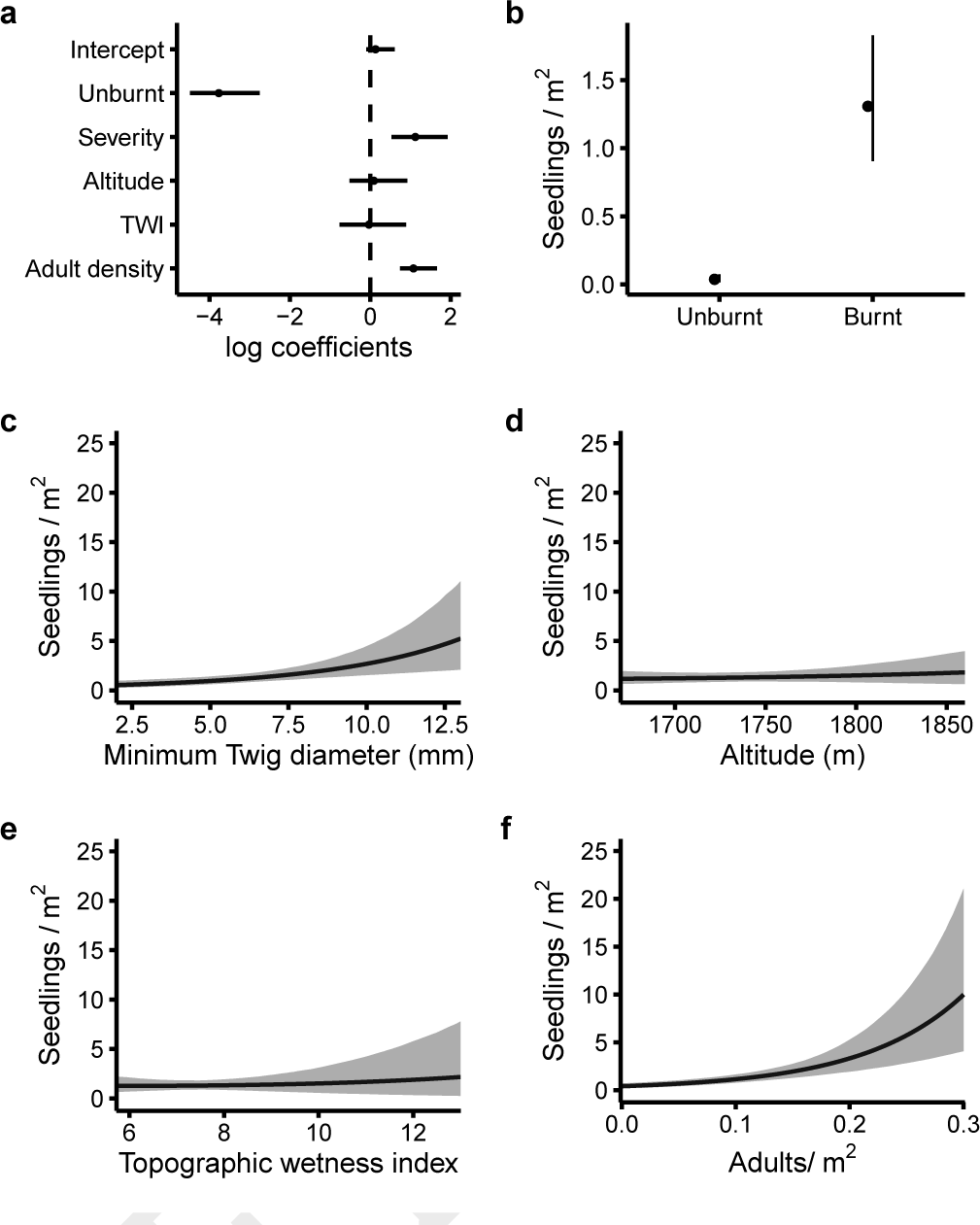
Mean *Grevillea australis* seedling density as a function of burning (burnt/unburnt), fire severity (twig diameter), altitude, Topographic Wetness Index (TWI) and adult density. (a) Centered and standardized model coefficients (on the log scale); and effects of: (b) burning, (c) fire severity, (d) altitude, (e) topographic wetness and (F) adult density, in areas burnt by the 2003 fires. All bars and shaded areas indicate 95% Bayesian Credible Intervals.

### Experimental warming and shrub seedling growth

Given that seedlings were more abundant in burnt vegetation, we investigated how warmer post-fire conditions affected seedling growth and mortality rates. Seedlings of *Grevillea, Asterolasia, Phebalium* and *Prostanthera* emerging after a wildfire were transplanted into experimentally burnt plots. These plots were either subjected to ambient conditions (i.e. controls), or enclosed in Open Top Chambers (OTCs) which increased temperature by 0.9°C.

After 2182 days (1281 growing season days) or 6 years’ growth, mean seedling heights of the tall shrubs (*Grevillea, Phebalium* and *Prostanthera*) growing in post-fire bare ground were greater in warmed plots relative to controls by 11.8, 4.5 and 14.9 cm, respectively (Fig. 3a). Warming increased heights of both *Grevillea* and *Prostanthera* seedlings in all years, while *Phebalium* did not respond to the warming treatment until the second growing season. By contrast, seedlings of the prostrate shrub, *Asterolasia*, showed no difference in growth rate between warmed and control plots in any year. For each species, similar treatment effects were observed for stem diameter growth (Fig. S4). Accounting for initial height and assuming logistic growth, the rates of change in mean annual predicted height of *Grevillea, Phebalium* and *Prostanthera* were 2.5, 1.6 and 2 times that observed in control plots, respectively. According to this model, a 6 cm seedling (the mean initial height of seedlings used in this experiment) attains maximum height 42 (*Grevillea*) or 18 years sooner (*Phebalium* and *Prostanthera*) when warmed by 0.9°C (Fig. 3b). *Asterolasia* was predicted to reach its maximum height in approximately 23 to 27 years, irrespective of warming treatment.

**Fig. 3.**
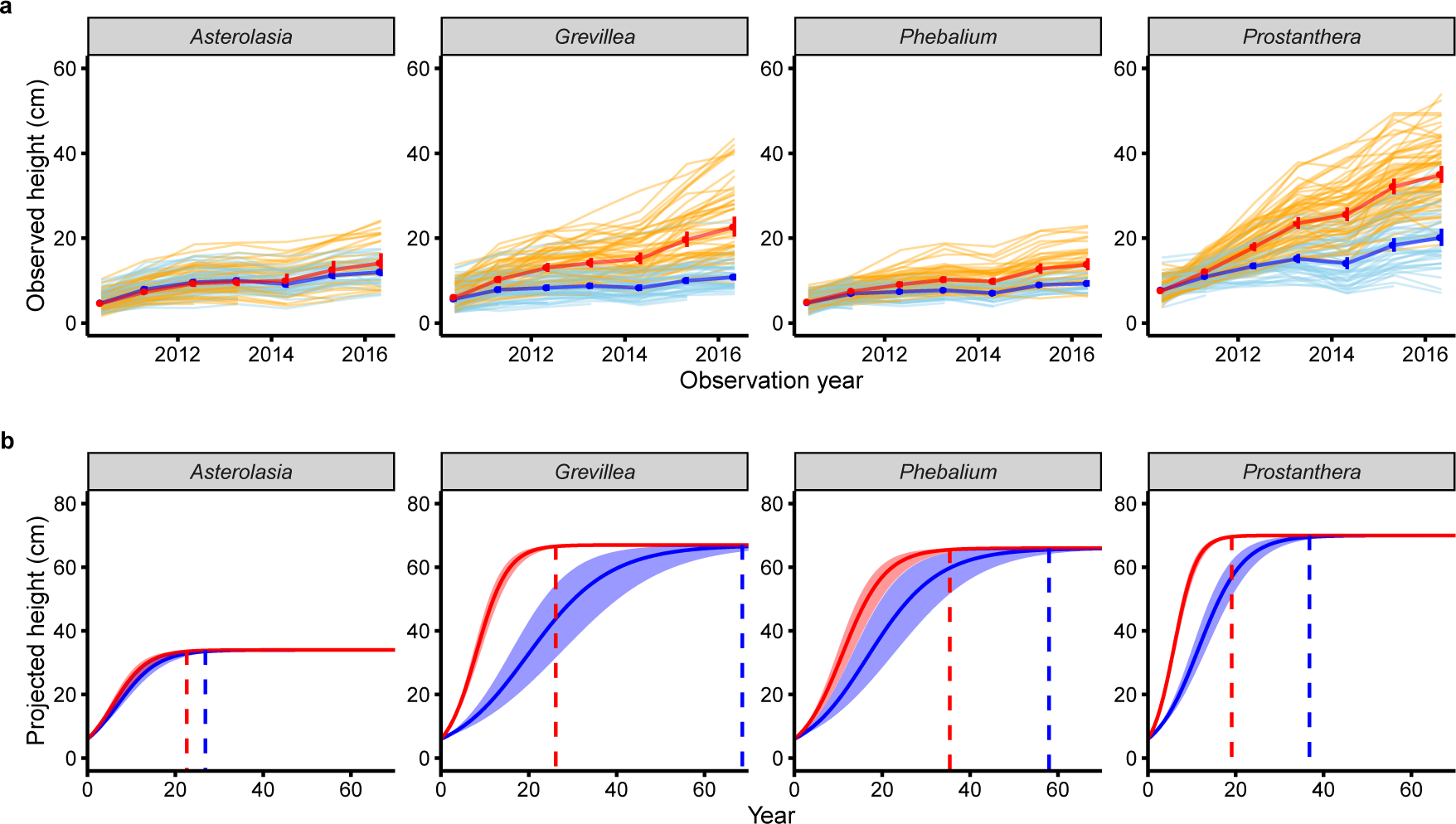
Observed and projected growth trajectories of four dominant Australian alpine shrubs. (a) Observed height growth: Thick lines with error bars represent mean (*±* 95% confidence intervals) observed heights at each May census. Thin lines represent individual growth trajectories. (b) Mean (*±* 95% Bayesian Credible Interval) projected growth trajectories. In all cases, red and orange lines = seedlings growing in warmed (OTC) conditions and blue and light blue lines = seedlings growing in control conditions. Projections were based on an logistic growth model using initial seedling size of 6 cm (the mean initial height observed in the OTC experiment) and mean maximum heights observed in long-unburnt sites (i.e. 34, 67, 66 & 70 cm for *Asterolasia, Grevillea, Phebalium*, & *Prostanthera*, respectively). Vertical lines delimit year in which maximum height is obtained

The growth responses of both *Grevillea* and *Asterolasia* observed in experimentally warmed plots were validated by the maximum heights of post-fire recruits across 30 open heathland sites burnt in 2003. Here, a 190 m altitudinal range is equivalent to a mean ambient temperature difference of approximately 1.5°C (Brown & Millner, 1989), which is comparable to that observed between experimentally warmed and control plots (0.9°C). In response to shifts in temperature, the maximum height of *Grevillea* post-fire recruits was expected to decrease with elevation, whereas *Asterolasia* seedlings were not expected to show this pattern. Our experimental predictions were verified (Fig. S5). Mean maximum height of *Grevillea* seedlings in burnt open heathland were 8 cm taller at 1670 m a.s.l compared to seedlings at 1860 m a.s.l. (22 cm vs 14 cm; a difference comparable to our experimental findings). In contrast, mean maximum height of *Asterolasia* seedlings did not vary significantly with elevation. Topographic Wetness Index and fire severity had no detectable influence on maximum seedling heights in either species.

### Experimental warming and shrub seedling mortality

After six years and across all plots, 36% (185 out of 511) of all seedlings transplanted into the 4×4 bare ground square grid had died. Most deaths occurred in *Asterolasia* (67) and *Phebalium* (67), followed by *Grevillea* (36) and *Prostanthera* (15). *Prostanthera* showed the largest treatment effect (Fig. 4), with annual mortality rates estimated to be near 0% in warmed plots and 4% in control plots. This significant decrease in mortality may be a consequence of OTCs reducing the severity of spring frosts by rising minimum ambient and soil temperatures by 1.1°C and 2°C, respectively (Fig. S6-S8). Warming also reduced mean seedling mortality in *Grevillea* and *Phebalium* (Fig. 4); however, for both species, the effect was highly uncertain (i.e. credible intervals overlap). By contrast, annual mortality rates in the prostrate shrub, *Asterolasia*, were marginally higher in warmed plots, but again this effect was highly uncertain (Fig. 4).

**Fig. 4.**
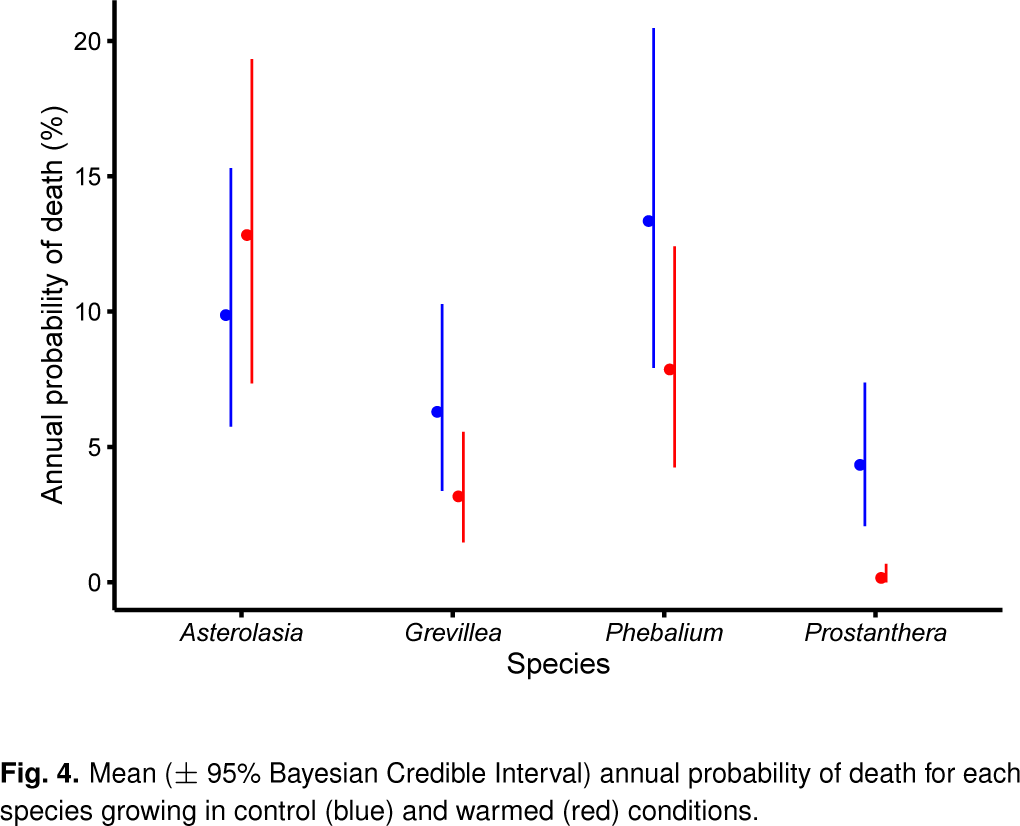
Mean (*±* 95% Bayesian Credible Interval) annual probability of death for each species growing in control (blue) and warmed (red) conditions.

### Effects of tussock grass proximity on shrub seedling growth and mortality rates

Climate change is expected to alter biotic interactions because their strength and direction depend strongly on climatic conditions, particularly in alpine and arctic ecosystems (Callaway *et al*., 2002; Klanderud, 2005). Here, we assess the interactive effects of warming and grass proximity on the growth and survival of *Grevillea* seedlings transplanted into various sized inter-tussock gaps. We detected a strong positive effect of warming treatment on growth rates and a marginally non-significant decrease in mortality (Fig. 5). However, we did not detect significant inter-tussock gap size effects or an interaction between gap size and warming treatment on either growth or mortality rates (i.e. coefficient credible intervals overlap zero).

**Fig. 5.**
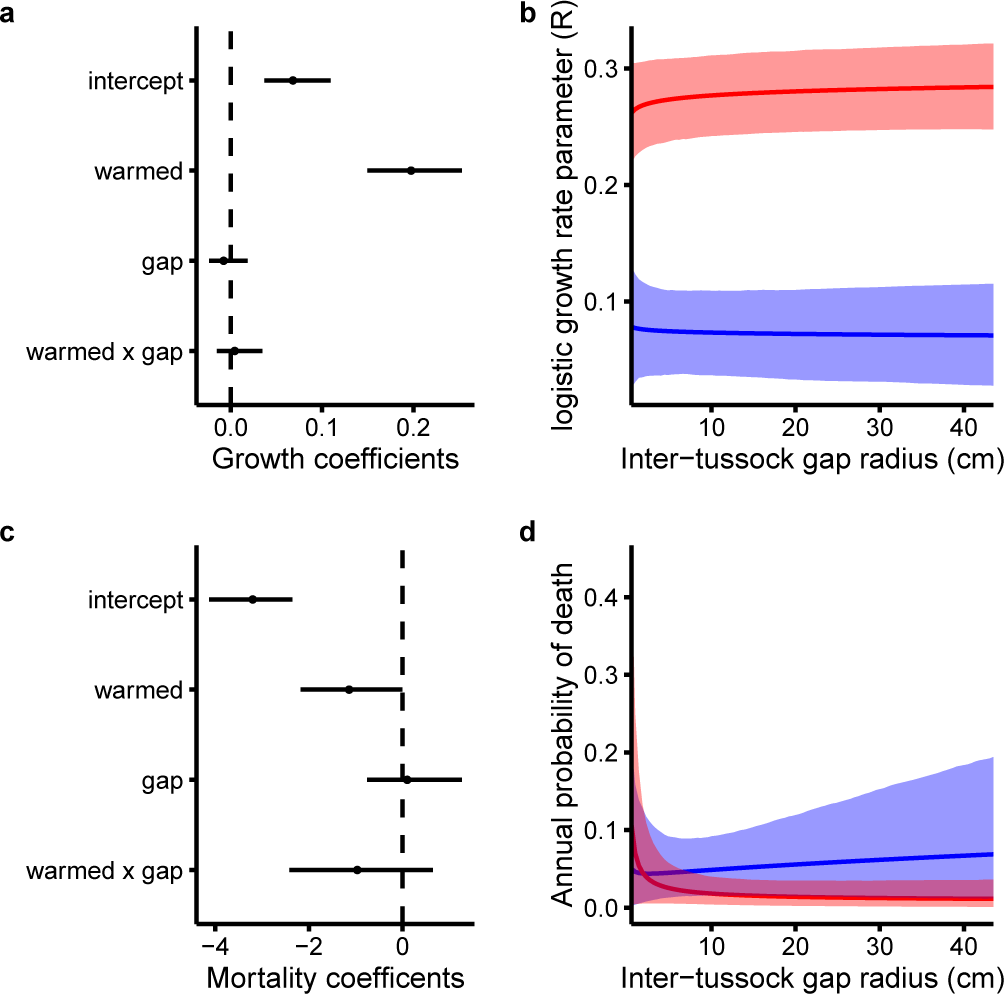
Experimental warming and inter-tussock gap size effects on *Grevillea australis* seedlings. Top rows = Growth rate effects, Second row = Mortality effects. (a & c) Centered and standardized model coefficients; (b & d) growth and mortality rate response curves along an inter-tussock gap size gradient in warmed (red) and control (blue) conditions. All error bars and shading are 95% Bayesian Credible Intervals.

### Experimental warming and rates of gap infilling by tussock grass

Despite having little impact on shrub seedling growth and mortality rates, tussock grass may still limit shrub recruitment, and thus, shrub expansion, by infilling post-fire bare ground gaps (whether by vegetative growth or seedlings) faster under warmer conditions. Using six years of post-fire inter-tussock gap size changes in warmed and unwarmed plots, we found that gaps were being infilled by tussock grasses (Fig. 6). However, the rate at which this occurred was very slow, with a 10 cm radius gap predicted to decrease by approximately 3 cm over a ten year period. We also detected no significant effect of a 0.9°C temperature rise on the rate of infilling.

**Fig. 6.**
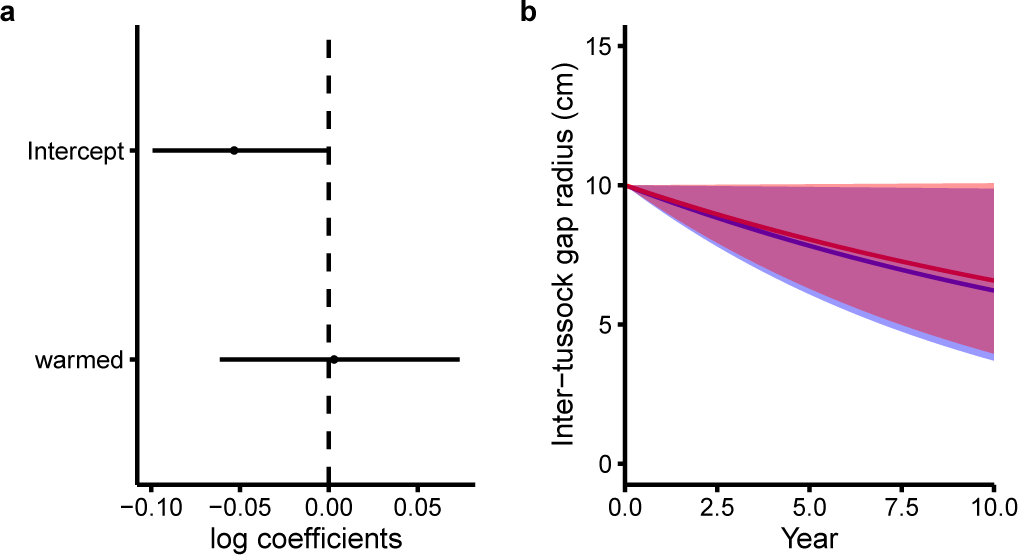
Effects of warming on rates of gap infilling by tussock grass. (a) Model coefficients and (b) projected temporal change in size for an average 10 cm inter-tussock gap in warmed (red) and unwarmed (blue) conditions. All error bars and shading are 95% Bayesian Credible Intervals.

## Discussion

In this study we have quantified several unknown interactions between fire, shrub-grass relationships and climate to extend a conceptual model of alpine shrub dynamics under climate change (Fig. 1b). In particular, we have shown how these interactions strengthen a hypothesized feedback that can rapidly increase shrub cover in the Australian Alps, and potentially, other alpine and tundra ecosystems (Higuera *et al*., 2008). While high cover of tussock grass is known to inhibit the establishment of shrub seedlings (Williams & Ashton, 1987), fire can create bare ground neccesary for shrub seedling establishment. Our results highlight that fire can increase shrub seedling establishment by as much 33–fold. Our results also indicate that if a shrub seedling manages to establish in a bare ground patch, its rate of growth and survival are not affected by tussock grass proximity, irrespective or warming. More importantly, our findings indicate that for tall shrubs, a 1°C increase in mean growing season temperature will result in a doubling of growth rates and a potential increase in survival. The consequence of these demographic effects will likely manifest as shrub thickening within and at shrub community boundaries. This in turn, will facilitate shrub establishment, via increased seed pools, into non-shrubby communities.

Landscape flammability, and thus fire, are also likely to increase as a result of warming effects on shrub seedling vital rates. Evidence from field studies of burning patterns (Williams *et al*., 2006), quantification of fuel mass and architecture (Fraser *et al*., 2016) and paleoecological studies (Higuera *et al*., 2008, 2009) indicate that shrubs are the most flammable component of alpine and tundra ecosystems (significantly more than tussock grassland; Fraser *et al*., 2016). In the Australian Alps, this differential flammability resulted in approximately 60% of open heathland and only 13% of tussock grasslands being burnt by the extensive 2003 wild-fires. Consequently, our results indicate that flammable fuel loads will accumulate twice as fast under a warmer climate. Ultimately this will further strengthen the feedback between shrubs and fire by increasing the frequency and severity of fires, which in turn, will create more bare ground, and thus more shrub recruitment opportunities both within and beyond current shrub boundaries —Recruitment opportunities that could persist for decades (Williams *et al*., 2014).

While we have addressed several unknowns between climatic warming, shrubs and fire, there are others we have not addressed that may also strengthen, weaken or break this feedback. The most obvious mechanism that will break this cycle involves short fire intervals that prevent fire-killed shrubs reaching reproductive age and thereby exhausting the seedbank (Enright *et al*., 2015). However, this scenario is unlikely for the majority of alpine (or tundra) landscapes, including those in Australia. For example, in the Australian Alps, current fire intervals of 50 to 100 years would need to decrease to less than 20 years—the time estimated for the species in this study to reach reproductive maturity (Williams *et al*., 2008). Furthermore, if reproductive output is related to plant size (Wenk & Falster, 2015), then climatic warming may allow fire-killed species to reach reproductive maturity sooner, and consequently, may increase their resilience to short fire intervals. Nevertheless, this and other factors such as changes in snow pack (Wipf *et al*., 2009), soil moisture (Engler *et al*., 2011), herbivory (Post & Pedersen, 2008) and adaptation (Byars *et al*., 2009; Hoffmann & Sgrò, 2011) require further research because they are all likely to be altered by the interactive effects of climate and disturbance in unpredictable ways.

By focusing on the life stage most vulnerable to climate and disturbance, and which determines a species capacity to establish in new areas (Walck *et al*., 2011; Briceño *et al*., 2015), our analyses provide a possible explanation as to why shrub cover is increasing in the Australian Alps, often at the expense of grasslands (McDougall, 2003; Wahren *et al*., 2013). An explanation that may also apply to other arctic and alpine ecosystems (Racine *et al*., 2004; Myers-Smith *et al*., 2011; Frost *et al*., 2013; Lantz *et al*., 2013). But more importantly, our results provide evidence for the underlying processes that could result in a warming-fire-shrub feedback that has been hypothesized in arctic paleoecological studies (Higuera *et al*., 2008, 2009). Based on current observations, average global temperature has already increased by 0.85°C since 1880 and is expected to rise by as much as 4.8°C by 2100 (IPCC, 2013). In alpine and tundra environments, temperatures (Chapin III *et al*., 2005), shrub cover (Myers-Smith *et al*., 2011) and the frequency and severity of fire (Westerling *et al*., 2006; Flannigan *et al*., 2009; Qiu, 2009; Bradstock *et al*., 2014) have all increased in the last few decades. These changes mean that the warming-shrub-fire feedback identified here is likely to have already strengthened. If this is the case, other non-woody communities will become shrubbier and more flammable, the effects of which, will have significant consequences for carbon sequestration, water supply and biodiversity.

## ACKNOWLEDGMENTS

This research was funded through Australian Research Council Linkage Grants, partnered through the Department of Sustainability and Environment, Parks Victoria, Commonwealth Scientific and Industrial Research Organisation (CSIRO). Plot infrastructure was supported by the Long Term Ecological Research Network. Funding was also received from the Centre of Excellence for Environmental Decisions (CEED) and Holsworth Research Committee. J.S.C was a recipient of an Australian Postgraduate Award. Monica Camac, Brad Farmilo, Monica Hursburgh, Paulius Kviecinskas, Karen Stott, Imogen Fraser, Sarah Bartlett, Paul Smart, Steven Rumbold, Lachlan Yourn and Chris Jones provided assistance with field measurements. William Morris and John Baumgartner provided coding advice and Warwick Papst provided field logistic advice. Daniel Falster and Richard Fitzjohn for developing software that enabled these analyses to be readily reproducible. Special thanks to John Baumgartner, David Bowman, Manuel Esperón-Rodríguez, Anaïs Gibert, John Morgan, Kimberley Millers, Michaela Plein and Inka Veltheim for their comments on this manuscript.

